# Multiple loci of small effect confer wide variability in efficiency and resistance rate of CRISPR gene drive

**DOI:** 10.1101/447615

**Authors:** Jackson Champer, Zhaoxin Wen, Anisha Luthra, Riona Reeves, Joan Chung, Chen Liu, Yoo Lim Lee, Jingxian Liu, Emily Yang, Philipp W. Messer, Andrew G. Clark

## Abstract

Gene drives could allow for control of vector-borne diseases by directly suppressing vector populations or spreading genetic payloads designed to reduce pathogen transmission. CRISPR homing gene drives work by cleaving wild-type alleles, which are then converted to drive alleles by homology-directed repair, increasing the frequency of the drive in a population. However, resistance alleles can form when end-joining repair takes place in lieu of homology-directed repair. Such alleles cannot be converted to drive alleles, which would halt the spread of a drive through a population. To investigate the effects of natural genetic variation on resistance formation, we developed a CRISPR homing gene drive in *Drosophila melanogaster* and crossed it into the genetically diverse *Drosophila* Genetic Reference Panel (DGRP) lines, measuring several performance parameters. Most strikingly, resistance allele formation post-fertilization in the early embryo ranged from 7% to 79% among lines and averaged 42±18%. We performed a Genome-Wide Association Study (GWAS) using our results in the DGRP lines and found that the resistance and conversion rates were polygenic, with several genetic polymorphisms showing relatively weak association. RNAi knockdown of several of these genes confirmed their effect, but their small effect sizes implies that their manipulation will yield only modest improvements to the efficacy of gene drives.

## INTRODUCTION

Super-Mendelian gene drive inheritance enables rapid spread of drive alleles in a population, even if these alleles cause harm to the organisms carrying them^1,2^. In homing drives, the drive allele contains an endonuclease that cleaves at a target site in the genome, and the drive allele is then copied into the target site during homology-directed repair, increasing the drive frequency in a population. A homing drive could thus facilitate the quick spread of a genetic payload, allowing a range of diverse applications. For example, a successful gene drive blocking disease transmission could be a powerful method to eliminate vector borne disease such as malaria, which kills nearly 500,000 people, mostly children, each year^3^. Gene drive constructs could also directly suppress a vector population by targeting an essential gene^1,2^. Other applications include the suppression of agricultural pest populations such as *Drosophila suzukii*^4^ or the removal of invasive species.

All these applications depend on functioning and highly efficient gene drives. However, there are practical barriers that reduce gene drive efficiency. One crucial barrier is the formation of resistance alleles against the drive allele^5–7^. With the development of CRISPR, homing drives have used Cas9 for their endonuclease component, which finds the target site with a guide RNA (gRNA). Cas9-based homing drives are highly susceptible to resistance because when Cas9 cleavage is repaired by end-joining instead of homology-directed repair, the target sequence is often changed, preventing recognition by the gRNA. Resistance has been found in all studies thus far involving *Drosophila*^6,8–12^, *Anopheles*^13–16^, and mice^17^. Recent efforts have made substantial progress against resistance alleles using multiplexed gRNAs^9^, improved promoters^9,13^, and careful choice of target sites to reduce formation of viable resistance sequences^13^, but it remains unclear if these improvements are sufficient for success in large cage populations, much less in highly diverse natural populations.

In most cases, resistance alleles form in the early embryo after cleavage by maternally deposited Cas9 and gRNAs. An early study indicates that genetically diverse lines experience high levels of variation in the rate at which these embryo resistance alleles form^6^. To further understand how genetic diversity may contribute to resistance allele formation, we tested a drive construct in the genetically diverse *Drosophila* Genetic Reference Panel (DGRP) lines. Several gene drive performance parameters were assessed, particularly the rate at which resistance alleles formed in the early embryo due to its wide range among lines. We were able to identify putative genetic factors affecting drive performance by conducting a genome-wide association study (GWAS) and further verified the results by knocking down candidate genes with lines expressing siRNAs and assessing the effect on drive performance.

## METHODS

### Genotypes and phenotypes

The CRISPR homing gene drive used for this study was constructed and assessed previously^6^. It carries a dsRed fluorescent protein gene driven by the 3xP3 promoter for expression in the eyes, ocelli, and the abdomen. The drive disrupts *yellow*, an X-linked gene, causing a recessive yellow body and wing phenotype. If Cas9 cleavage is repaired by end-joining, rather than homology-directed repair, this will mutate the target site, forming a resistance allele. These are termed “r2” if they render *yellow* nonfunctional due to a frameshift or a sufficient change in the amino acid sequence. Resistance alleles that preserve the function of *yellow* are termed ‘r1’. The different possible phenotypes and genotypes are summarized in the SI Dataset, as are calculations based on phenotypes to determine drive performance parameters. A similar drive targeting *white*^9^ was also assessed in three DGRP lines.

### Fly rearing and phenotyping

All flies were reared on Bloomington Standard medium at 25°C with a 14/10 hr day/night cycle. During phenotyping, flies were anesthetized with CO_2_ and examined with a stereo dissecting microscope. Flies were considered ‘mosaic’ if any discernible expression of yellow phenotype could be observed in the body or wings. dsRed fluorescent phenotype in the eyes was scored using the NIGHTSEA system (SFA-GR). All experiments involving live gene drive flies were carried out using Arthropod Containment Level 2 protocols at the Sarkaria Arthropod Research Laboratory at Cornell University, a quarantine facility constructed to comply with containment standards developed by USDA APHIS. Additional safety protocols regarding insect handling approved by the Institutional Biosafety Committee at Cornell University were strictly obeyed throughout the study, further minimizing the risk of accidental release of transgenic flies.

### Fly experiments

To assess drive performance in the DGRP lines, males with the drive were crossed to virgin females from the DGRP lines. The progeny, which consisted of females heterozygous for the drive and wild type males, were allowed to mate and then placed into 4-6 vials. These flies were removed after five days. Progeny of these crosses were collected after eleven, fourteen, and seventeen days. They were frozen and later scored for dsRed and yellow body phenotype, as well as white eye phenotype, which occurred in approximately half of males inheriting the original drive allele.

To assess the effect of siRNAs on drive performance, females homozygous for the homing drive were crossed separately to *w*^-^/Y;GMR-Gal4/*CyO* males and *w*^-^/Y;Act-Gal4,*w*^+^/*CyO* males. Females from the former cross with *CyO* were then crossed to males of the latter cross without *CyO*. Male progeny with dsRed, yellow body, wild type eyes, and *CyO* from this cross with were then crossed with siRNA and control lines. These included lines 36303, 36304, 38263, 40849, 41875, 51417, 51435, 51933, 52882, 61942, and 67932 from the Bloomington *Drosophila* Stock Center and lines 60100 and 104478 from the Vienna *Drosophila* Resource Center. Female progeny from these crosses with wild type eyes, wild type body, and without *CyO* were then crossed to *w*^1118^ males, and the progeny were scored for dsRed, white eye phenotype, and yellow body phenotype.

### Genome-wide association study

DGRP genotype files were downloaded from the DGRP2 website (http://dgrp2.gnets.ncsu.edu/data.html). 128 lines from these files were phenotyped, and drive performance parameters were calculated (Data S1-S2) for GWAS analyses. To compensate for the cryptic relatedness among DGRP lines, PLINK 1.9^18^ and filters of minor allele frequency (maf 0.05) and genetic information missingness (geno 0) were used to modify the genotype file and to generate a genetic relatedness matrix using GEMMA^19^. Then filters (maf 0.05 and geno 0.05) were applied to the genotype of the 128 lines. Phenotype data were added to the genotype file as new columns. The *Wolbachia* infection status was obtained from the DGRP2 website and evaluated as a covariate in the analysis.

Association tests were performed using GEMMA and the following univariate linear mixed model adapted from a study by Zhou and Stephens^19^:

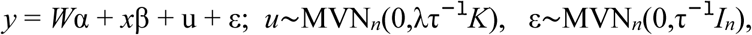

where *y* is an *n*-vector of quantitative traits for *n* strains, *W*is a matrix of covariate *Wolbachia*, α is the intercept, *x* is an *n*-vector of marker genotypes, and β is the effect size of the marker. *u* is an *n*-vector of random effects with an *n*-dimensional multivariate normal distribution (MVN_*n*_) that depends on λ, the ratio between the two variance components, τ^−1^, the variance of the residual errors, and *K*, the known *n* × *n* relatedness matrix. ε is an *n*-vector of residual errors with a MVN_*n*_ that depends on τ^−1^ and *I_n_*, the identity matrix.

A Wald test was performed and resulting test statistics were generated and loaded into R studio to build Manhattan plots. To compensate for family-wise error rate, *P*-values were adjusted using the BH function^20,21^. To verify that the cryptic relatedness among DGRP lines was properly compensated for, we used the R package “qqman” (http://www.gettinggeneticsdone.com/2014/05/qqman-r-package-for-qq-and-manhattan-plots-for-gwas-results.html) to draw Q-Q (quantile-quantile) plots. The genomic inflation factor λ was calculated by converting the observed and expected *P*-values to χ^2^ statistics and taking the ratio of the median of the observed χ^2^ to the median of the expected χ^2^ Top hits were manually annotated using FlyBase^22^.

## RESULTS

### Drive characteristics vary widely among the DGRP lines

Males with a CRISPR homing drive targeting the X-linked *yellow* gene were crossed to females from 128 DGRP lines, each of which has a conserved target site sequence. The resulting offspring (drive heterozygote females and wild-type males) were allowed to mate, and their progeny were phenotyped for dsRed and yellow body and wings to assess drive performance (Data S1-S2). Germline drive conversion rate varied among the DGRP lines, with a range of line means from 0.282 to 0.717, and averaging 52±9% (Figure S1, Figure 1A). This among-line variation was found to be highly significant (*P* < 2×10^−16^, *F*=3.422, ANOVA). Resistance allele formation was also measured. These can take the form of r2 alleles, which disrupt *yellow*, and r1 alleles, which preserve *yellow* function. Based on male progeny, the average rate at which r2 resistance alleles were formed was 38±8% (Figure S1), and the remaining 7±4% either had r1 resistance alleles formed or remained wild-type (Figure 1B). From female progeny data, the rate at which r2 resistance alleles formed in the early embryo in flies that inherited the drive had a wide range of line means (from 0.072 to 0. 794; *P*<2×10^−16^, *F*=22.26, ANOVA), averaging 42±18% (Figure S1, Figure 1C). In addition to r2 resistance alleles that form in the early embryo and give a full yellow phenotype, slightly later cleavage can result in a mosaic phenotype (Figure 1D). The level of mosaicism is related to the rate of the full embryo resistance alleles (Figure S2A). Thus, we examined the difference between the observed rate of mosaicism and the expected level and found moderate variation among the DGRP lines (Figure S2B).

**Figure 1.**
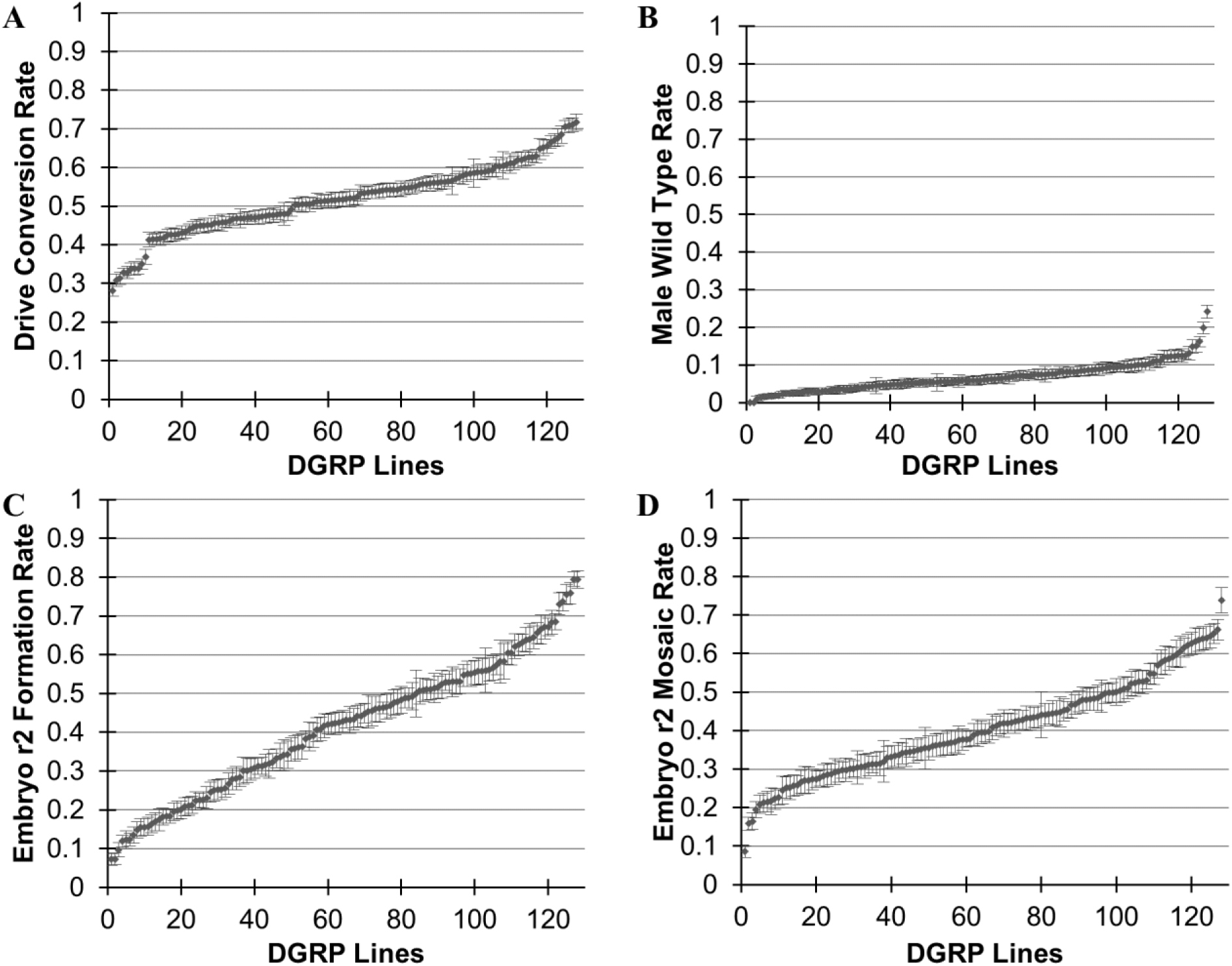
Variation in drive performance among the DGRP lines. Each point represents the rate ± standard error in a single line for **(A)** drive conversion efficiency in heterozygote females, **(B)** the rate of wild-type phenotype in male progeny, **(C)** the rate of full embryo r2 resistance allele formation, and **(D)** the rate of embryo mosaicism.

Examining three DGRP lines with low, medium, and high embryo resistance with a separate gene drive targeting the *white* gene with otherwise identical components produced a similar pattern of embryo resistance (Data S3, Figure S3), indicating that much of the variation observed is independent of target site.

### Genome-wide association study identifies genes affecting drive performance

Our DGRP phenotype data allowed us to identify genetic variants that affect the gene drive performance and assess their effect sizes. A total of 903,282 single nucleotide polymorphisms (SNPs) were included in the cryptic relatedness-compensated genome-wide association study using the phenotype data from our 128 DGRP lines. A univariate linear mixed model was applied, and the Wald test was performed to estimate a *P*-value for each variant. To compensate for family-wise error rate (FWER), *P*-values were adjusted using the Benjamini-Hochberg approach. Adjusted *P*-values were ranked in ascending order. One compelling result from this study is that despite the large and highly significant genetic variance among lines in the scored drive phenotypes, no marker SNP attainted genome-wide significance after multiple-testing correction. This implies that rather than having large effect variants in the genetic background, the variation was highly polygenic. Even with a 50% (false discovery rate), the drive conversion (Figure S4A) and germline r2 resistance from male data phenotypes produced no hits.

For the germline wild-type rates in the male data, several significant hits were obtained.However, despite a λ of 1.016, the Q-Q plot illustrates that the *P*-values were severely inflated because the phenotypic distribution showed low variance and right-skewness (Figure 1B, Figure S4B). The seemingly normal genomic inflation factor is due to the fact that the median of the test statistic was not affected much by the hits with the lowest 1% of *P*-values. However, these hits were the most inflated, as shown by a high λ of 1.110 for the top 10,000 hits. As GWAS generally requires normally distributed data, we hypothesized that the inflation was due to the data structure, and we performed rank normalization to the data. After normalization, the Q-Q plot showed less inflation (Figure S4B) (λ=1.033 for the entire data, and λ=1.014 for the top 10,000 SNPs), and no hits were obtained using a false discovery rate of 50%.

For early embryo r2 resistance allele formation rate, 63 polymorphisms were found using a very relaxed 50% false discovery rate threshold (Data S1, Figure 2A). A Q-Q plot was generated (Figure S4C), and the genomic inflation factor of λ=1.052 showed that *P*-values were not substantially inflated due to relatedness or other factors. The top hits were annotated for location relative to known genes, and the functions of such genes were also manually annotated. Functions most likely to be considered related to the embryo r2 rate were transcription and translation factors, mRNA and protein degradation factors, genes related to DNA repair, and gene of unknown function. Preferred location was within a gene or a potential promoter region, but not deep within an intron (within 1 kb of a coding sequence). Based on these locations and functions, a selection of most promising genes was obtained, including *CG5009*, *Dlish*, *CG7220*, *Camta*, and *pum* (Table 1). The top two hits in the analysis were in the *pyd* gene, but since they were deep within an intron of this gene, it was not included in the list of most promising genes.

**Figure 2.**
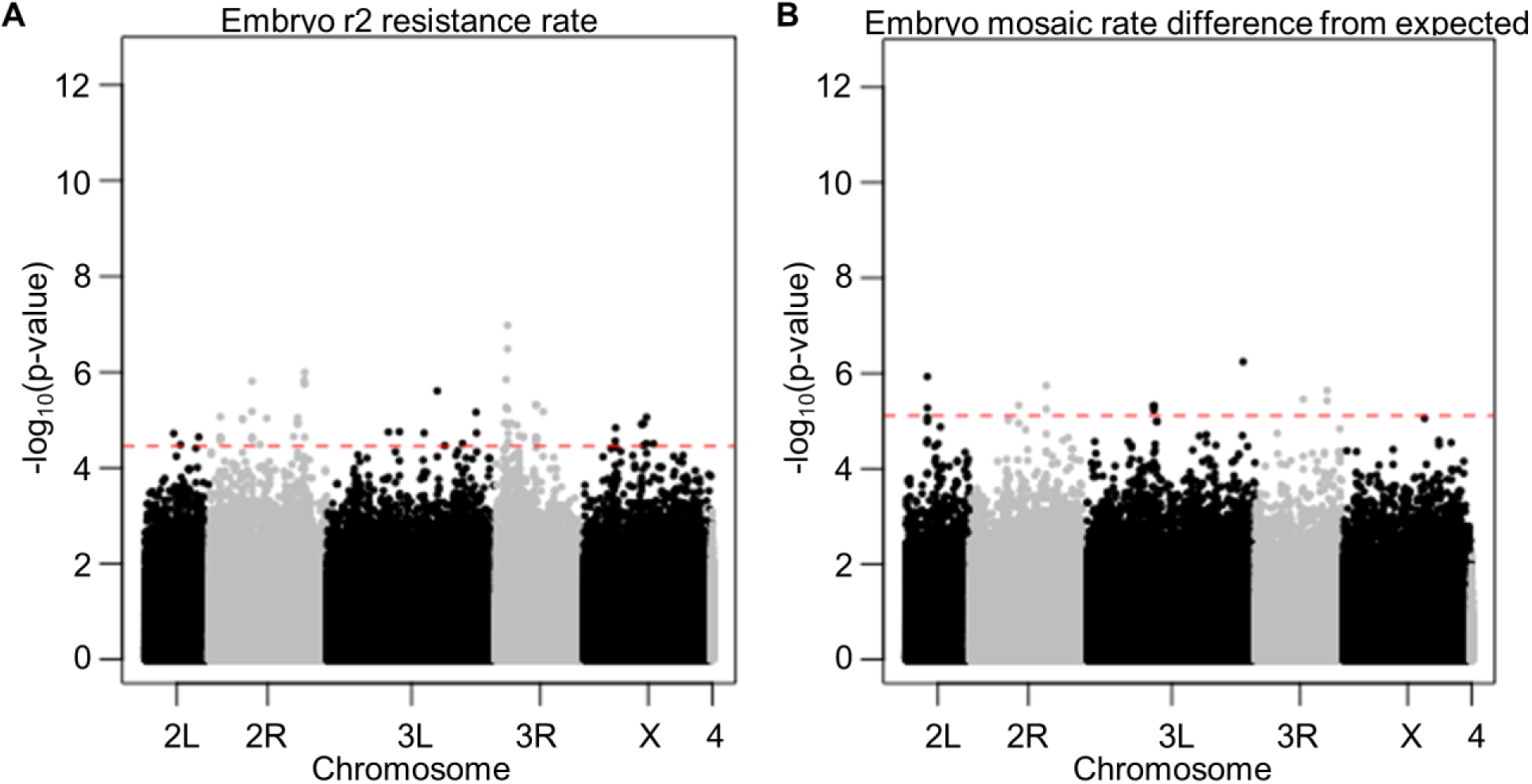
Manhattan plot shows top hits from GWAS analysis. Each dot shows the location and *P*-value of a single polymorphism of the **(A)** early embryo r2 resistance allele formation rate and **(B)** embryo mosaic rate difference from the expected value. The red dashed line shows the *P*-value cutoff corresponding to a 50% false discovery rate.

**Table 1.** Top GWAS hits for embryo resistance rates

| SNP location | $P$-value | Hits | Gene | Location | Molecular function |
| --- | --- | --- | --- | --- | --- |
| **Full early embryo r2 resistance allele formation rate** |  |  |  |  |  |
| chr2R:17762194 | 1.01E-06 | 2 | *CG5009* | coding sequence | dehydrogenase/oxidase |
| chr2R:17723899 | 1.48E-06 | 1 | *Dlish* | intron | cadherin binding |
| chr2R:10738229 | 1.54E-06 | 4 | *CG7220* | 3'UTR | Ubiquitin-conjugating |
| chr2R:9457770 | 9.58E-06 | 2 | *Camta* | promoter/intron | transcription activator |
| chr3R:9081589 | 1.27E-05 | 3 | *pum* | intron | post-transcriptional repressor |
| chr3R:12693319 | N/A | N/A | *Arp1* | coding sequence | cytoskeleton/microtubule |
| chr3R:12657943 | N/A | N/A | *CG6225* | coding sequence | aminopeptidase |
| **Embryo mosaic rate difference from the expected value** |  |  |  |  |  |
| chr2R:15324753 | 1.81E-06 | 2 | *Arf51F* | 3'UTR | GTP hydrolase |

Note: only the hit with the lowest $P$ -value for each gene is shown. N/A indicates top hits from an earlier analysis with fewer DGRP lines, but not providing any significant hits in the full analysis.

In addition to the full embryo r2 resistance allele formation rate, we also analyzed the difference between the mosaicism rate compared to the expected mosaicism rate based on the embryo r2 rate (Figure 2B, Figure S4D). This adjustment allowed us to factor out the polymorphisms associated with the embryo r2 rate and focus on those that allowed Cas9 activity to persist for particularly long or short periods of time beyond the early embryo. Twelve hits were obtained with a 50% false discovery rate (Data S5), of which a single gene, *Arf51F*, was considered a top hit based on location and function (Table 1).

### siRNA knockdown of GWAS hits affects resistance allele formation

*D. melanogaster* lines expressing short hairpin RNA (shRNA) under control of the UAS promoter against the genes from Table 1 were obtained, together with control lines, to verify our top GWAS hits. These were introgressed into drive heterozygote females together with a GAL4 gene under control of the beta-actin promoter. These flies were crossed to *w*^1118^ males, and the resulting progeny were phenotyped to assess drive performance (Data S6). After multiple testing correction, siRNAs against *Camta* was found to significantly reduce the rate of embryo r2 resistance formation from 42% to 26%, while siRNA against *pum* increased the embryo resistance rate to 60% (Figure 3).

**Figure 3.**
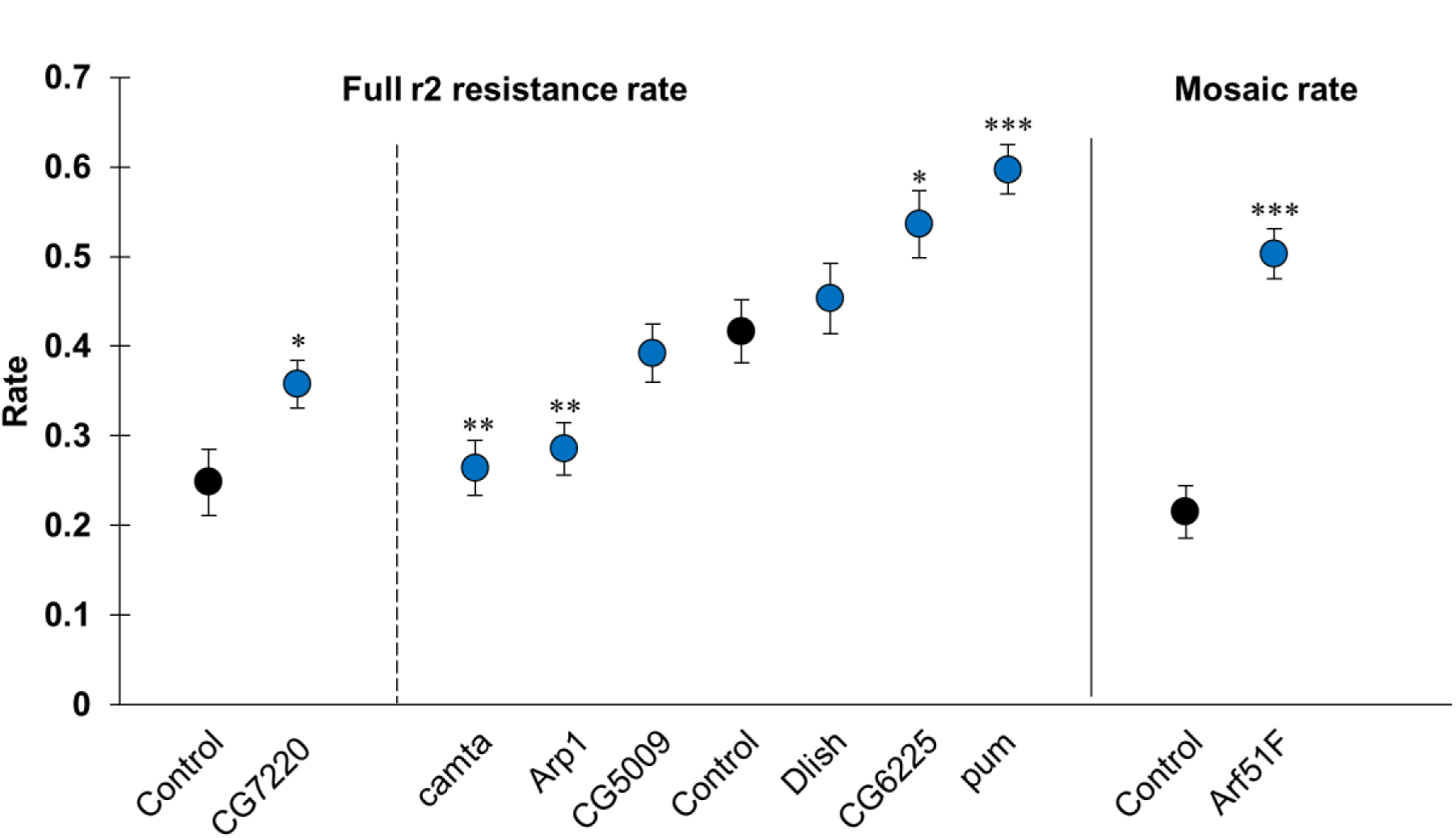
siRNAs against hits from GWAS affect the embryo resistance rate. Each point represents the embryo resistance rate ± standard error in a single shRNA line. The dashed line separates two sets of shRNA lines that each correspond to a different control line. The solid line separates lines analyzed for full embryo r2 resistance and mosaic resistance. \**P*<0.05, \*\**P*<0.01, \*\*\**P*<0.001, Fisher’s exact test, comparing to corresponding control line. After multiple testing correction, only the marked ** or *** were considered significant.

Two more genes, *Arp1* and *CG6225* (Table 1), were also included in the shRNA analysis (Figure 3). These were top hits from a preliminary analysis, but not our full analysis including all DGRP lines tested. Of these, *Arp1* had a statistically significant effect, reducing the embryo resistance rate from 42% to 29%. Additional shRNAs were tested against *Lig4* and *Irbp* (Ku70), genes known to be involved in end-joining pathways and therefore, possibly involved in resistance allele formation. However, no significant effects were observed after multiple-testing correction on any performance parameters (Data S6).

An siRNA against *Arf51F* was found to increase the embryo r2 mosaicism rate from 22% in the control line to 50% (Data S6, Figure 3), which was statistically significant (*P*<0.001, Fisher’s exact test). The difference in the full embryo resistance rates between these lines was less than 1%. Note also that no statically significant results were observed when assessing the differences in the mosaicism rates between shRNA lines tested for the full embryo r2 phenotype.

## DISCUSSION

Here, we examined natural variation in the performance of CRISPR homing gene drives. We found that the rate of resistance allele formation in the embryo varies widely among lines, indicating that there is naturally-occurring genetic variation affecting the efficacy of gene drives. This is particularly noteworthy, since embryo resistance is a primary obstacle gene drives must overcome to attain performance sufficient for success in the wild^6,9,14–16^. A GWAS analysis covering 128 DGRP lines did not reveal any genetic loci of large effect size for embryo resistance rate, and as is often the case with these studies, no marker attained a genome-wide false discovery rate less than 5%. This is of high significance for application of gene drives in natural populations, as it implies that *D. melanogaster* has extensive polygenic variation (i.e. many loci of small effect) that can potentially respond to selection to increase resistance to gene drives. The GWAS with greatly relaxed stringency (false discovery rate of 50%) did identify several putative gene target hits after extensive curation, which were tested for effect by shRNA-based knockdown. This revealed several genes affecting embryo resistance and may provide some insight into relevant pathways that may be the cause of evolved resistance. However, the effects sizes of the variants in the DGRP lines are uniformly small, implying that these genes would not likely be highly useful targets for manipulation to improve drive performance.

We expect embryo resistance to be caused by maternally deposited Cas9 protein and RNA, in addition to gRNA. Thus, factors that could directly affect this include transcription and translation levels, in addition to the degradation rates of the RNAs and Cas9 protein. Our GWAS and siRNA verification revealed several genes with functions that could potentially affect these, including *camta*, a transcription factor, and *pum*, a post-transcriptional repressor. The potential effect of *Apr1*, which is involved in the cytoskeleton or microtubules, on embryo resistance allele formation is less clear. *Arf51F*, a GTP hydrolase, also appeared to affect the rate at which Cas9 protein or mRNA can persist into later stages of the embryo and form mosaic resistance alleles, though it apparently did not affect the rate in the early embryo. Since mosaic individuals could still perform successful drive conversion according to a previous study ^12^, it is unclear how it may affect overall drive performance, though presumably population suppression drives or population replacement drives targeting haplolethal genes may be negatively affected by additional mortality or sterility in individuals inheriting the drive.

The results presented here beg the question of how gene drives might be better engineered for improved efficiency or to be unresponsive to the background genetic variation that we detected. Since some of the siRNAs, namely *camta* and *Arp1*, reduced the rate of early embryo resistance allele formation, it may be possible in future studies to reduce resistance rates by lowering expression of these genes. These siRNAs could potentially be used together, multiplexed with several gRNAs using the tRNA^23–25^ and/or ribozyme^26–29^ systems, which also have more flexibility in their promoter. Indeed, the U6 promoters most often used for gRNAs may not be appropriate for shRNA use, since these shRNAs may impose a moderate or even severe fitness cost. This is likely, since natural variants in the DGRP lines did not affect the phenotype to the same degree as the shRNA knockdown, implying that a similar level of knockdown is selected against. However, even if an improved drive construct incorporating such an RNAi gene functioned successfully, our data indicates that embryo resistance would only be modestly reduced. The polygenic basis of the variation in resistance that was quantified here poses one of the most serious challenges to perfection of an effective gene drive for deployment in a natural population. Though we only tested this in *D. melanogaster*, there is no reason to expect that other gene drive systems would be much different.

Our results underscore the need to test gene drives in a set of genetically diverse lines in the specific target species to gain a more realistic understanding of how they may perform in natural populations. Among the DGRP lines, we found that individuals could vary more than 10-fold in embryo resistance, an effect that would be further exaggerated in natural populations compared to our experiments, in which the drive heterozygotes all shared 50% of their genome, with variation only occurring in the 50% from the DGRP lines. Modeling efforts simulating the release of gene drives should also take such variation into account, since certain individuals or regions with particular polymorphisms may experience much higher or lower rates of resistance.

## ACKNOWLEDGEMENTS

Thanks to Yassi Hafezi for helpful advice on the experimental work, Robert Unckless for useful discussions and comments on the study, and Ana Jaksic for helpful comments on the manuscript. This study was supported by startup funds from the College of Agriculture and Life Sciences at Cornell University to P.W.M, the National Institutes of Health award R21AI130635 to J.C., A.G.C., and P.W.M, and the National Institutes of Health award F32AI138476 to J.C.

---

**Figure S1.**
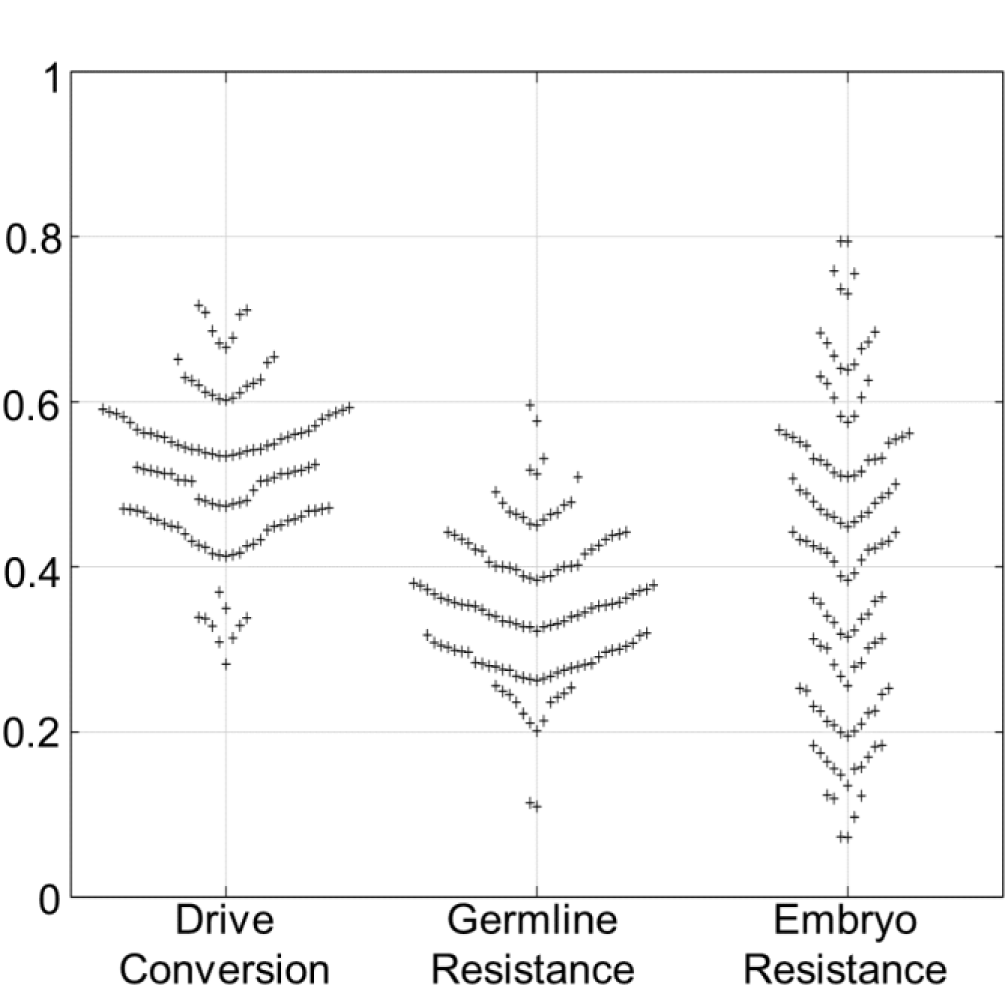
Variation in drive performance parameters among the DGRP lines. Each symbol on the plot represents the average rate for a single DGRP line.

**Figure S2.**
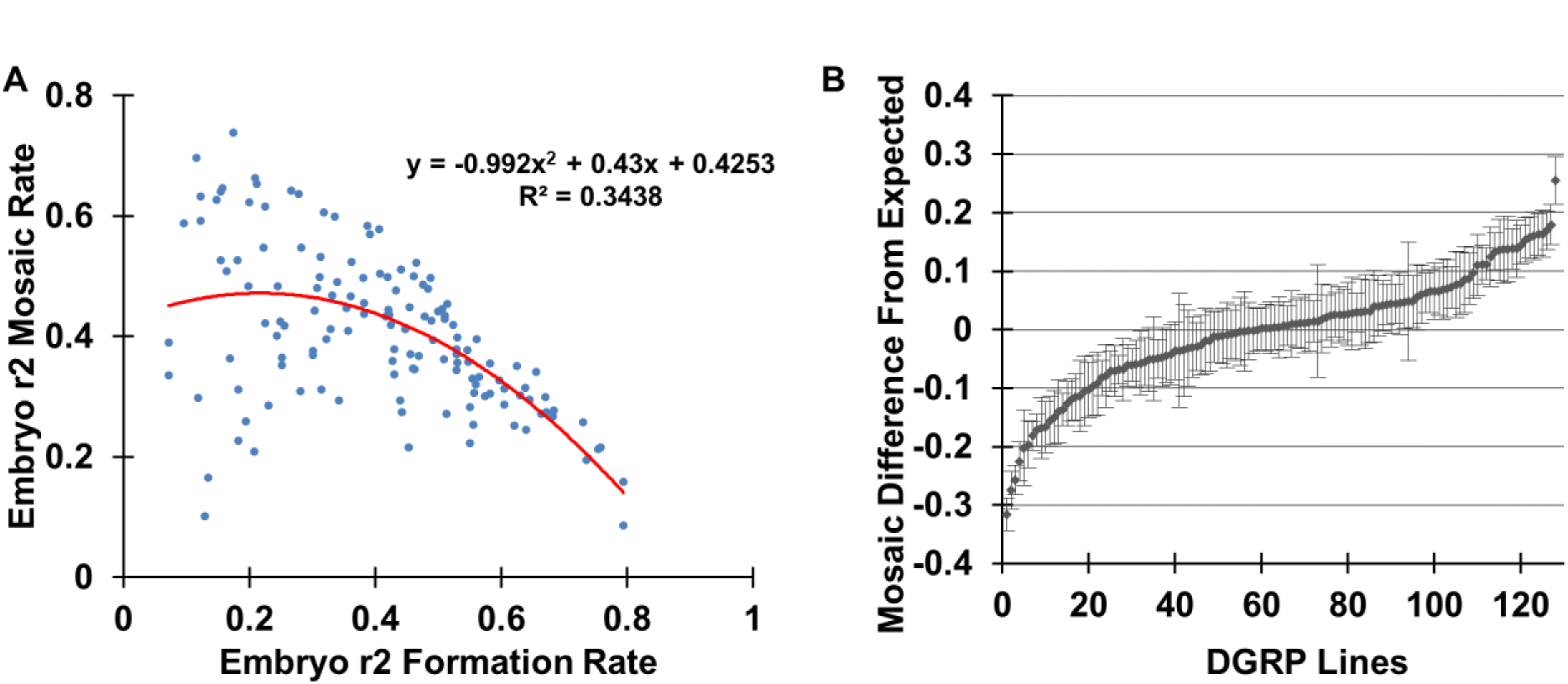
The relation between the full embryo r2 resistance allele formation rate and the mosaic embryo rate. **(A)** Each point represents values from a single DGRP line. A second-order polynomial was fitted to the data to determine the average expected level of mosaic individuals based on the level of individuals with full r2 alleles. **(B)** The difference between the observed and the expected level of mosaicism was plotted. Each point represents the difference value ± standard error in a single DGRP line.

**Figure S3.**
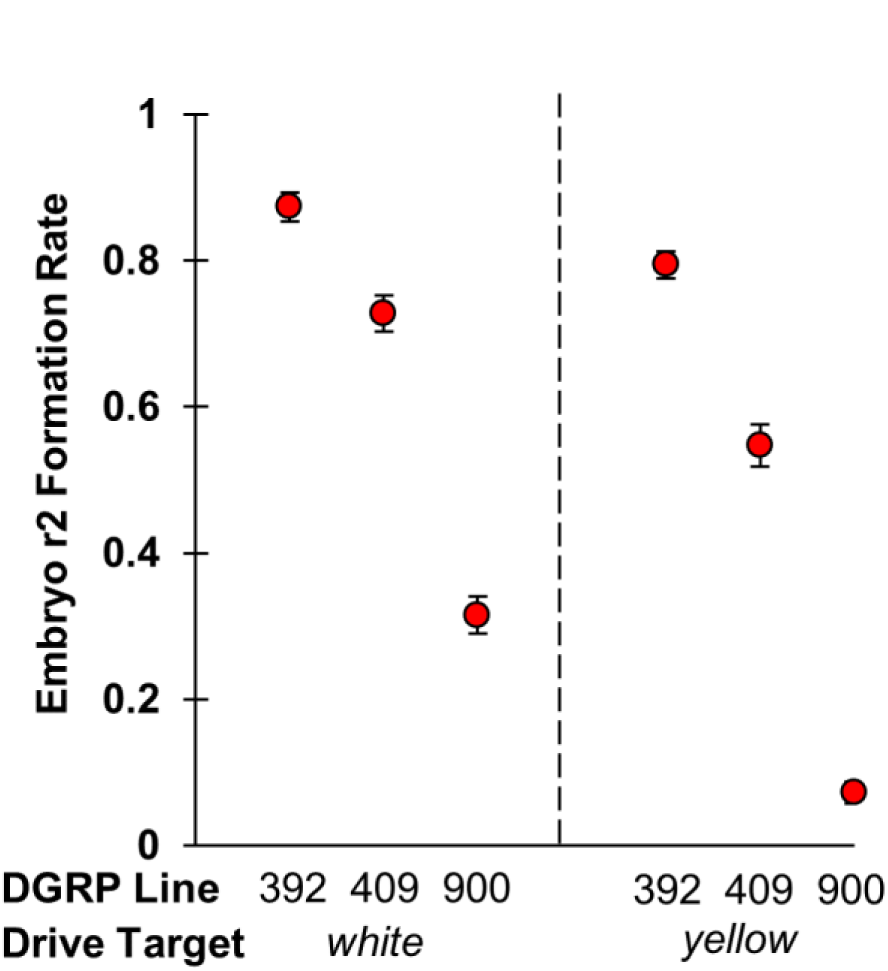
Effects of genetic background are conversed between drives with different target sites. The rate of full embryo r2 resistance allele formation is shown (± standard error) for three DGRP lines with both the drive targeting *yellow* and a similar drive targeting *white*. The drive targeting *white* produces overall more resistance allele formation, but the pattern between the lines is similar.

**Figure S4.**
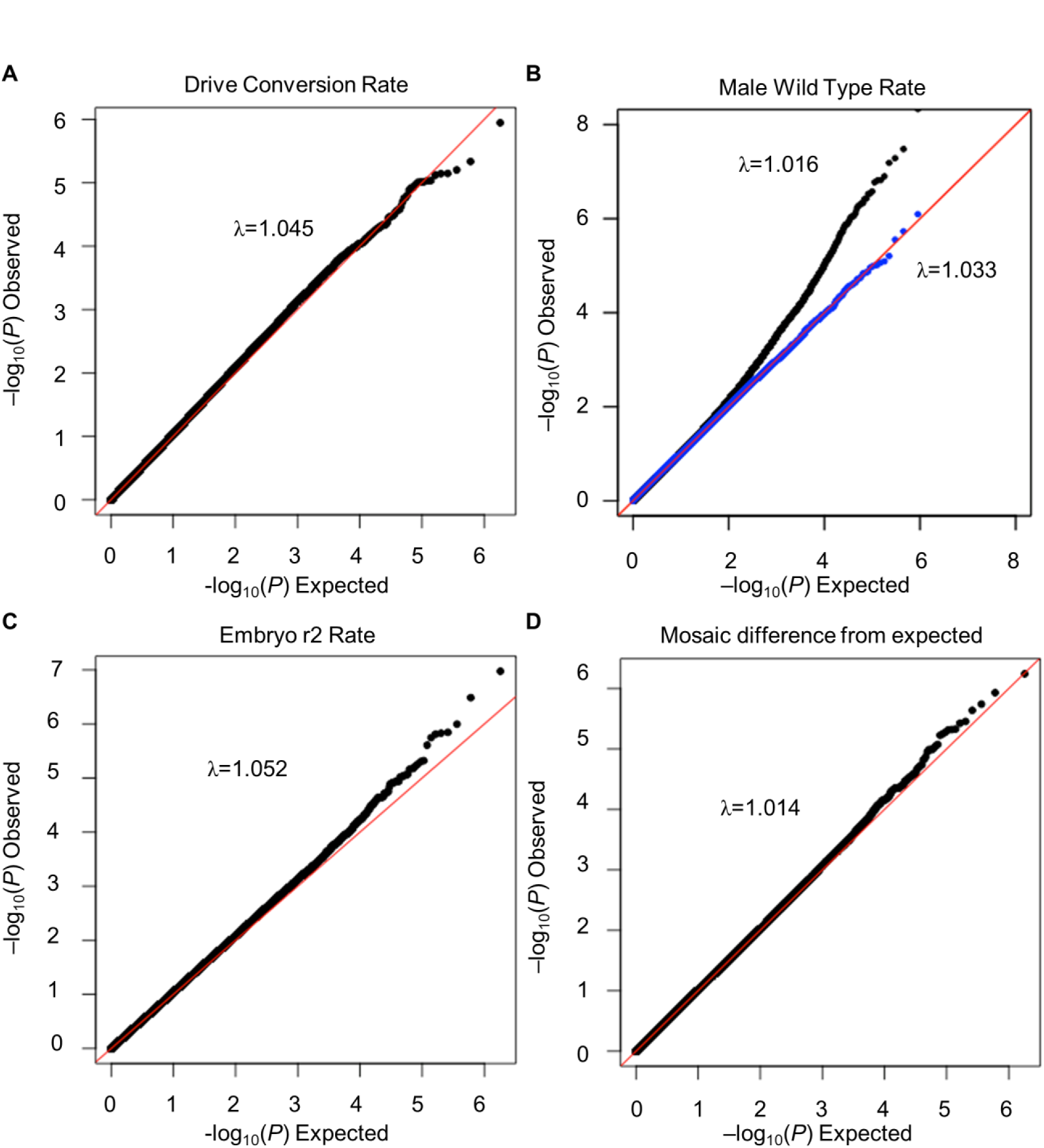
Q-Q analysis of GWAS for drive performance among the DGRP lines. A plot of expected vs. observed −log_10_(*P*-value) for each hit for **(A)** drive conversion efficiency in heterozygote females, **(B)** the rate of wild-type phenotype in male progeny, with rank-normalization corrected values shown in blue, **(C)** the rate of full embryo r2 resistance allele formation, and **(D)** the difference between expected and observed embryo mosaic rate.

**Figure S5.**
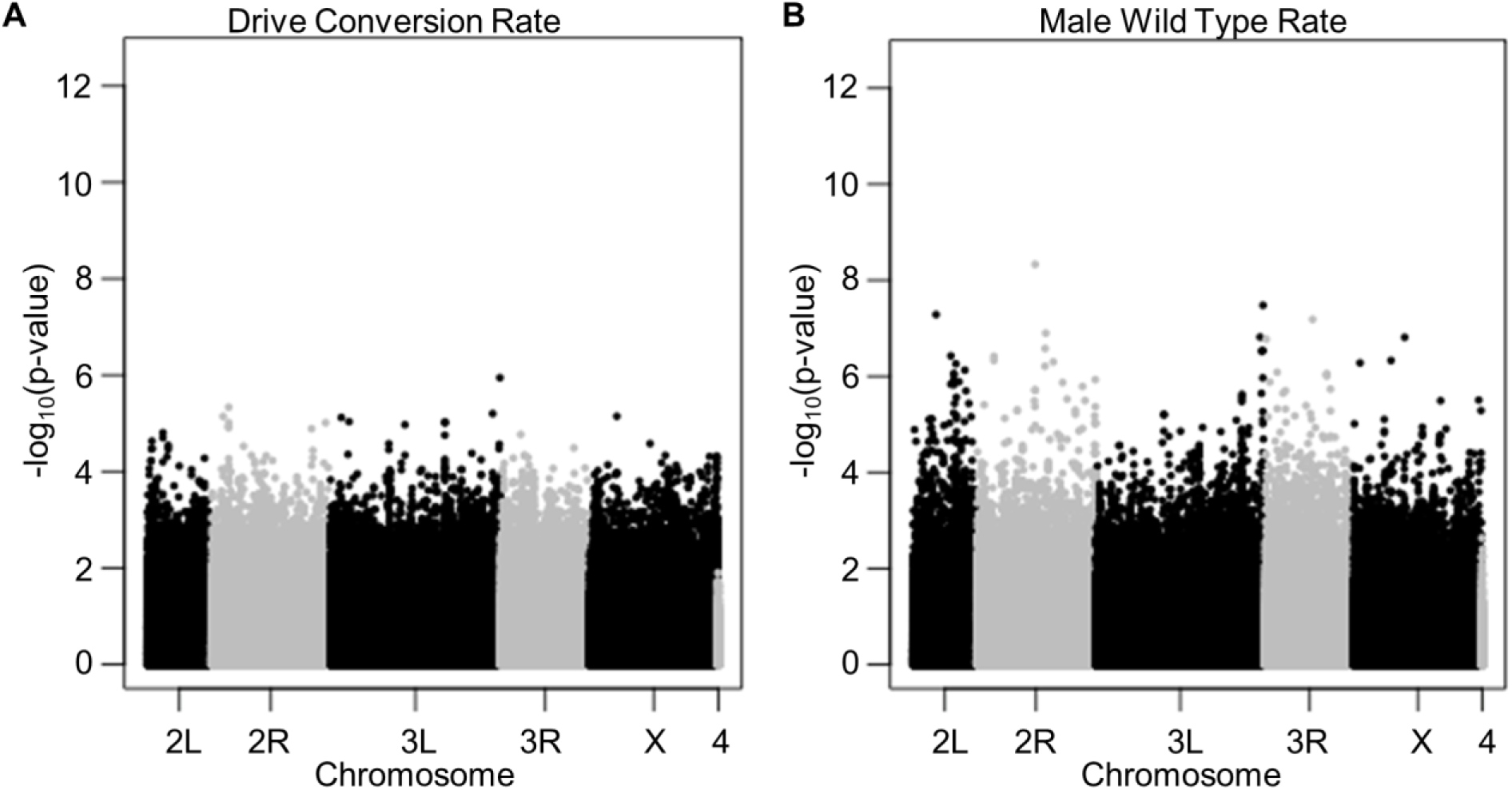
Manhattan plot from GWAS analysis. Each dot shows the location and *P*-value of a single polymorphism of the **(A)** drive conversion rate in heterozygote females and **(B)** the rate of wild-type phenotype in male progeny.

